# Enforced specialization fosters mutual cheating and not division of labour in the bacterium *Pseudomonas aeruginosa*

**DOI:** 10.1101/2021.06.26.450018

**Authors:** Subham Mridha, Rolf Kümmerli

**Affiliations:** Department of Quantitative Biomedicine, Winterthurerstrasse 190, 8057 Zurich, Switzerland

**Keywords:** Public goods, Siderophores, Bacterial communities, Iron availability, Negative frequency dependent fitness, Regulatory networks

## Abstract

A common way for bacteria to cooperate is via the secretion of beneficial public goods (proteases, siderophores, biosurfactants) that can be shared among individuals in a group. Bacteria often simultaneously deploy multiple public goods with complementary functions. This raises the question whether natural selection could favour division of labour where subpopulations or species specialise in the production of a single public good, whilst sharing the complementary goods at the group level. Here we use an experimental system, where we genetically enforce specialization in the bacterium *Pseudomonas aeruginosa* with regard to the production of its two siderophores, pyochelin and pyoverdine, and explore the conditions under which specialization can lead to division of labour. When growing pyochelin and pyoverdine specialists at different mixing ratios in various iron limited environments, we found that specialists could only successfully complement each other in environments with moderate iron limitation and grow as good as the generalist wildtype but not better. Under more stringent iron limitation, the dynamics in specialist communities was characterized by mutual cheating and with higher proportions of pyochelin producers greatly compromising group productivity. Nonetheless, specialist communities remained stable through negative frequency-dependent selection. Our work shows that specialization in a bacterial community can be spurred by mutual cheating and does not necessarily result in beneficial division of labour. We propose that natural selection might favour fine-tuned regulatory mechanisms in generalists over division of labour because the former enables generalists to remain flexible and adequately adjust public good investments in fluctuating environments.

## Introduction

Division of labour is a defining feature of major transitions in evolution (Szathmáry and Smith, 1995; Ispolatov *et al*., 2012; Rueffler *et al*., 2012; Bourke, 2011). At all levels of biological organization, division of labour underlies specialization, whereby a generalist entity transits to multiple specialist entities, performing complementary tasks. Examples include the transition from (i) generalist RNA molecules serving as genetic code and enzymes to specialized DNA (genetic code) and proteins (enzymes) within cells, (ii) generalist cells to specialized soma and germlines in multicellular organisms, and (iii) generalist individuals to specialized castes in eusocial insect societies (Bourke, 2011; Cooper and West, 2018). Division of labour entails that specialized individuals (or entities) perform cooperative tasks that benefit others in the group and that specialization leads to an inclusive fitness benefit to all individuals (West and Cooper, 2016). Consequently, division of labour is more likely to evolve and remain stable when the interests of the specialized individuals are aligned, which is typically the case when relatedness is high and the traits individuals specialize on are essential for survival and reproduction (Fisher *et al*., 2013; West and Cooper, 2016).

Despite this clear conceptual framework, it is often difficult to experimentally demonstrate that a particular form of specialization constitutes division of labour. Due to their high amenability, bacterial (and other microbial) systems have become popular models to test basic theory and to experimentally investigate division of labour and the underlying evolutionary forces (Ackermann *et al*., 2008; van Gestel *et al*., 2015; Kim *et al*., 2016; West and Cooper, 2016; Dragoš *et al*., 2018a; Dragoš *et al*., 2018b; Giri *et al*., 2019; Armbruster *et al*., 2020; Yanni *et al*., 2020). Bacteria are social organisms, and the most common form of cooperation involves the secretion of beneficial compounds that can be shared as public goods (West *et al*., 2007). For example, bacteria secrete enzymes to digest extra-cellular polymers and proteins, quorum-sensing molecules for communication, biosurfactants to enable swarming on wet surfaces, and siderophores to scavenge iron (Drescher *et al*., 2014; Roman *et al*., 2017; Kramer *et al*., 2020; Yan *et al*., 2019). Because multiple public goods might be required at the same time, a key question is whether bacterial populations can segregate into subpopulations, each specializing in one of the public goods and mutually sharing the beneficial compounds at the population level (Frank, 2013; West and Cooper, 2016; Schiessl *et al*., 2019). Indeed, the evolution of such division of labour has been described in several laboratory systems. For example, *Bacillus subtilis* populations segregate in biosurfactant-producing and matrix-producing cell types which synergistically interact to enable sliding motility (van Gestel *et al*., 2015). Similarly, populations of *Pseudomonas fluorescens* rapidly segregate into two cell types, one producing a lubricant at the front of the colony and the other one pushing cells forward (Kim *et al*., 2016). Other studies enforced specialisation through genetic engineering to study its fitness consequences. In *Bacillus subtilis*, enforced specialization between two strains producing either the structural protein TasA or the exopolysaccharide EPS lead to stable division of labour that outperformed the generalist wildtype in forming pellicle biofilms (Dragoš *et al*., 2018a). Another study showed that a poor-performing engineered cross-feeding system between two *Escherichia coli* strains evolved into a productive, mutually beneficial division of labour during experimental evolution (Preussger *et al*., 2020).

In contrast to these positive results, there are also examples where division of labour was either not stable or did not evolve (Dragoš *et al*., 2018b; Schiessl *et al*., 2019). For example, a follow-up study on the TasA vs. EPS specialization in *B. subtilis* showed that division of labour collapsed through the evolutionary reversion to an autonomous lifestyle (Dragoš *et al*., 2018b). Moreover, a comparative study on hundreds of *Pseudomonas* isolates from natural pond and soil communities revealed high variation in public goods production among isolates, but little evidence for specialisation and division of labour (Kramer *et al*., 2020). Finally, a common phenomenon linked to public goods production is cheating, where mutants stop contributing to public goods production, yet exploit the public goods produced by others (Özkaya *et al*., 2017; Smith and Schuster, 2019). Cooperators and cheaters often co-exist in populations, which reflects a form of specialization (Maclean *et al*., 2010; Driscoll *et al*., 2011), but does often not represent division of labour because cheaters typically compromise group productivity (West and Cooper, 2016). The conditions that tip the balance in favour of division of labour as opposed to cheating are often unclear, but likely involve intrinsic (genetic architecture) and extrinsic (environmental) factors, determining the costs and benefits of specialization.

In our study, we tackle this issue by using a model system where we experimentally enforce specialization between two bacterial strains and measure its fitness consequences across a range of environmental conditions. Specifically, we worked with two engineered mutants of the bacterium *Pseudomonas aeruginosa*, each specialized in the production of one of its two siderophores, pyoverdine or pyochelin (Serino *et al*., 1997; Visca *et al*., 2007; Youard *et al*., 2011; Schalk *et al*., 2020). Siderophores are produced in response to iron limitation, to scavenge this essential nutrient from the environment (Kramer *et al*., 2020). Both pyoverdine and pyochelin can be public goods and shared between cells (Ross-Gillespie *et al*., 2015) and although they are simultaneously produced, they take on slightly different functions (Dumas *et al*., 2013). Due to its relatively low iron affinity, pyochelin is primarily beneficial and increasingly produced under moderate iron limitation, while pyoverdine has a very high iron affinity and is more relevant and produced in high amounts under severe iron limitation (Dumas *et al*., 2013; Ross-Gillespie *et al*., 2015; Mridha & Kümmerli, 2021).

Using this experimental system, we tested whether enforced specialization alleviates the metabolic burden associated with the simultaneous production of both siderophores (intrinsic factor), whilst the benefits of the two molecules can still accrue to all individuals through molecule sharing at the population level. To this end, we mixed the pyochelin and pyoverdine specialists across a range of mixing ratios and compared the growth of the entire population relative to the fitness of the generalist wildtype strain producing both siderophores. If the two strains engage in division of labour, we expect population growth to be higher for specialized mixes than for generalist cultures. Conversely, if the two specialists engage in mutual cheating, then we expect population growth of mixed cultures to be compromised relative to generalist cultures. Moreover, we tested whether extrinsic environmental factors influence the propensity of specialization leading to division of labour or mutual cheating by manipulating the iron availability in the medium. We predict that the potential for division of labour is highest at intermediate iron availability where both siderophores are beneficial, whereas division of labour should not occur under high iron availability (where siderophores are no longer required) and is less likely to occur under severe iron limitation, as pyochelin is less useful. Finally, we explored whether the two specialists can co-exist or whether one type drives the other one to extinction. For this purpose, we calculate the relative fitness of the two strains in co-culture across a range of mixing ratios. Co-existence would result in negative-frequency dependent fitness patterns, where both specialists win the competition when initially rare in the population and consequently settle at an equilibrium frequency.

## Materials and methods

### Bacterial strains

We used *P. aeruginosa* PAO1 (ATCC 15692) as our wildtype strain, which produces the siderophores pyoverdine and pyochelin. Additionally, we used its reporter variants where either a green (*gfpmut3*) or a red (*mcherry*) fluorescent gene was chromosomally integrated into the wildtype genetic background. For simplicity of nomenclature, the green fluorescent gene *gfpmut3* will be henceforth referred to as *gfp*. To investigate the effect of individual siderophores on growth, we used the deletion mutants producing only a single siderophore in the PAO1 strain background, namely, (1) *PAO1DpvdD*: a strain only producing pyochelin as the gene coding for the pyoverdine synthetase PvdD is deleted, and (2) *PAO1DpchEF*: a strain only producing pyoverdine as the gene coding for the pyochelin synthetases PchE and PchF are deleted. To determine the frequency of the two specialist siderophore producers in mixed cultures, we used the following variants with chromosomally integrated reporters (same genes as described for the wildtype above): (a) PAO1*DpvdD::gfp*, (b) PAO1*DpvdD::mcherry*, (c) PAO1*DpchEF::gfp*, (d) PAO1*DpchEF::mcherry*. All reporter constructs were integrated at the *att*Tn7 site using the mini-Tn7 system (Choi and Schweizer, 2006).

### Growth conditions

Prior to experiments, we prepared overnight cultures from −80 °C stocks, in 8ml Lysogeny broth (LB) in 50ml tubes, incubated at 37°C and 220 rpm for approximately 18 hours. Cells were harvested by centrifugation (8000 rpm for 2 minutes), subsequently washed in 0.8% saline, and adjusted to OD_600_=1 (optical density at 600nm). The harvested cells were grown in CAA medium (5g casamino acids, 1.18g K_2_HPO_4_*3H_2_O, 0.25g MgSO_4_*7H_2_O, per litre), buffered at physiological pH by the addition of 25mM HEPES. To induce a gradient of iron limitation, we used either plain CAA or CAA supplemented with 100μM and 300μM of the synthetic iron chelator 2-2’-bipyridyl. We further created an iron-replete condition by supplementing CAA with 100μM FeCl_3_. All chemicals were purchased from Sigma Aldrich (Buchs SG, Switzerland).

### Productivity of specialist and wildtype cultures

We first investigated whether mixtures of the two specialized siderophore producing strains grow better or worse than the generalist wildtype strain producing both siderophores. We mixed the strains PAO1Δ*pvdD::gfp* producing only pyochelin and the strain PAO1Δ*pchEF::mcherry* producing only pyoverdine at different starting frequencies in all the four media, differing in their iron availability. We used the following pyochelin:pyoverdine specialist mixing ratios (in %): 100:0, 95:5, 75:25, 50:50, 25:75, 5:95, 0:100, whereby 100% means specialist monocultures. We quantified the growth of these mixed and monocultures in parallel to the wildtype PAO1 monoculture. Experiments were carried out in 200μl medium distributed on a 96-well plate. Bacteria were inoculated at a starting density of OD_600_ = 0.0001 and incubated at 37°C for 24 hours in a multimode plate reader (Tecan, Männedorf, Switzerland). Cultures were shaken every 15 minutes for 15 seconds before measuring growth (OD_600_). We applied blank subtractions to account for the optical density of the media. We conducted four independent experiments with 2 to 3 replicates for each mixing ratio and media condition.

We used the OD_600_ as a proxy for bacterial biomass and considered the growth integral (area under the bacterial growth curve) over 24 hours as a measure of productivity. Because we were interested in the relative performance of mixed- and mono-cultures of specialists compared to the wildtype, we divided the growth integral of the specialist cultures by the growth integral of the wildtype monoculture for the respective media condition. Values of productivity > 1 or productivity < 1 indicate that specialist cultures perform better or worse than wildtype cultures, respectively.

### Measuring strain frequency with flow cytometry

For all mixed cultures of specialists, we measured the frequency of the fluorescence-tagged strains PAO1Δ*pvdD::gfp* producing only pyochelin and PAO1Δ*pchEF::mcherry* producing only pyoverdine before and after the growth experiments described above. Frequencies were measured with the LSRII Fortessa cell analyser (BD Biosciences, Allschwil, Switzerland) at the Flow Cytometry Facility of the University of Zurich. The GFP tagged cells (pyochelin producers) were recorded in the FITC channel (excitation = 495 nm, emission = 519 nm) using a blue laser at 488nm and 530/30 bandpass filter. The mCherry tagged cells (pyoverdine producers) were recorded in the PE-Texas Red channel (excitation = 566 nm, emission = 616 nm) using the yellow-green laser at 561nm and 610/20 bandpass filter. The Cytometry Setup and Tracking, settings of the instrument were used and the threshold for bacterial cell detection was set to 200V (lowest value possible).

Bacterial mixes at the start of the experiment were diluted in sterile 1x phosphate buffer saline (PBS; Gibco, ThermoFisher, Zurich, Switzerland) and 50,000 events were recorded at a low flow rate in the flow cytometer, yielding the initial (t_0_) frequency of the two siderophore specialists. After the 24 hours growth cycle in the plate reader (see section above), we processed the bacterial cultures to quantify the frequencies of fluorescently tagged specialists after competition (t_24_). We diluted the cultures in a 96-well plate, with 1x PBS depending on the endpoint optical density obtained from the plate reader. Then we recorded 50,000 events from 10 μl of these diluted cultures using a high-throughput sampler device (BD Bioscience) at a low flow rate.

To analyse the flow cytometry data, we used the software FlowJo (BD Biosciences, Ashland, USA). We followed a three-step gating strategy (Fig. S1): (a) we separated bacterial cells from debris and background noise based on the forward (area) and side scatter (granularity) values typical for bacterial cells. (b) Within that cell cluster we imposed the PE-Texas Red-H and forward scatter gates to distinguish between mCherry positive (pyoverdine producers) and negative cells. (c) Within the mCherry negative cell cluster, we then imposed the FITC-H and forward scatter gates to determine the GFP positive (pyochelin producers) cells. This stratified gating strategy enabled us to eliminate debris, noise, and dead cells from our analysis. We used the following three controls: (i) sterile 1x PBS to quantify the background noise of the medium; (ii) the untagged wildtype strain monoculture as a negative control to determine the gating threshold for non-fluorescent cells; (iii) the fluorescently tagged monocultures of specialists and the wildtype PAO1 strains as positive controls to determine the gating threshold for fluorescent cells.

### Relative fitness measurement

With the initial (t_0_) and final (t_24_) frequencies of the siderophore specialists, we calculated the relative fitness (*v*) of the pyochelin producer (PAO1Δ*pvdD::gfp*) following Ross-Gillespie *et al*. 2007 (Ross-Gillespie *et al*., 2007).

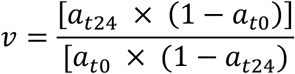

Here a_t0_ and a_t24_ are the frequencies of the pyochelin producer at the beginning and the end of the experiment, respectively. We subsequently log-transformed the relative fitness values. Values of ln(*v*) > 0 or ln(*v*) < 0 indicate whether the frequency of the pyochelin producer increased (i.e., won the competition) or decreased (i.e., lost the competition) relative to the pyoverdine producer.

We conducted a control experiment to test whether the two different fluorescent reporters affect strain fitness due to the implied metabolic costs. For this we competed identical strains of specialised siderophore producers against each other that only differ in their fluorescent reporters (i.e. PAO1Δ*pvdD::gfp* versus PAO1Δ*pvdD::mcherry* and PAO1Δ*pchEF::gfp* versus PAO1Δ*pchEF::mcherry*. We mixed both strain pairs in a 1:1 ratio and subjected them to the same four media conditions (differing in their level of iron limitation) as in the main experiments. We observed that strains expressing GFP experienced a fitness advantage (Fig. S2). This fitness advantage was relatively minor in iron-rich medium, yet gradually increased with more stringent iron limitations. To correct for this marker-based fitness bias, we calculated the average relative fitness of both GFP expressing strains in the control experiment for each media condition separately (Fig. S2, Table S1) and subtracted these values from the respective relative fitness values of PAO1*DpvdD::gfp* in our main experiment, where this strain competed against PAO1*DpchEF::mcherry*.

### Expected frequency estimation

If strains in mixtures do not socially interact then we expect that their fitness is determined by their intrinsic growth rates shown in monocultures. In contrast, if there are social interactions between strains – e.g., through the exchange of siderophores – then we expect that their relative fitness in mixtures significantly deviates from the relative fitness predicted from their intrinsic growth rates in monocultures. To estimate the expected frequency of strains in the absence of social interactions, we applied the procedure by Sathe & Kümmerli 2020 (Sathe and Kümmerli, 2020). In brief, we used the growth kinetic data of the monocultures *PAO1*Δ*pvdD::gfp* and PAO1*DpchEF::mcherry* to fit parametric growth curves (Gompertz function yielding the best fit). From these curve fits, we extracted the maximal growth rate μ, and estimated the time strains spent in the exponential phase t_exp_. The starting point is defined by the end of the lag phase (from the model fit), while the end point is defined by the entry to the stationary phase (upon visual inspection of the model fits using the image analysis software FIJI). Using μ and t_exp_, we then estimated the number of cell divisions per unit time as *k* = *μ*/ln2 and the number of generations as *n* = *kt*_exp_, for each strain and media condition. We then calculated the expected absolute frequency of both specialists after competition as *F*_end_ = 2^n^ x *F*_start_ and estimated the relative expected frequency of the pyochelin producer as *f*_pyochelin_ = *F*_end_pyochelin_ / (*F*_end_pyochelin_ + *F*_end_pyoverdine_). We then compared these expected relative frequencies to the actual relative frequencies observed.

### Statistical analysis

We used general linear models for statistical analysis in R 3.4.2. To analyse if siderophore specialists grow better or worse than the generalist wildtype we used analysis of variance (ANOVA) models with strain background and iron availability as fixed factors. To account for multiple pairwise comparisons between the two specialists and the wildtype we corrected p-values using the false discovery rate method. To test if the productivity of the specialist mixes varied with the initial frequency of pyochelin producers in the mixes and with iron availability, we used ANCOVA models with the initial frequency as covariate and iron availability as fixed factor. Subsequently we performed one-way ANOVA for each media separately and conducted two-sided one sample *t*-tests to determine whether productivity differs between specialist mixes and the generalist wildtype, which equals one. To investigate whether observed final frequencies of the pyochelin specialist differed than the expected frequencies, we used ANCOVA models, with frequency type (expected versus observed) and iron availability as fixed factors and initial frequency of pyochelin producers as covariate. Finally, we tested whether the relative fitness of the pyochelin producers depended on its initial frequency in the mixes. For this, we used ANCOVA models with initial frequency as covariate and iron availability as fixed factor, subsequently performing one-way ANOVA for each media separately.

## Results

### Pyoverdine-producing specialists grow better than pyochelin-producing specialists

To assess how the two specialist strains (pyochelin and pyoverdine producers) and the wildtype strain (producing both siderophores) perform in monoculture, we monitored their growth in four variants of the CAA media differing in their level of iron limitation. We found that productivity (growth integral) differed significantly across strains (Fig 1, Table S2, ANOVA: F_2,88_= 5.434, p = 0.0059) and decreased as the medium became more iron limited (F_3,88_= 736.9, p < 0.0001), while there was no interaction between the two variables (F_6,82_= 1.136, p = 0.3489). Specifically, we found that pyoverdine-producing specialists grew significantly better than pyochelin producing specialists (post-hoc comparison: p = 0.0046), whereas there were no significant differences between the productivity of the wildtype and the two specialists (post-hoc comparisons, wildtype vs. pyoverdine specialist: p = 0.2067; wildtype vs. pyochelin specialist: p = 0.0690). Overall, these results reveal that pyoverdine specialists outperform pyochelin specialist in monoculture, showing that pyoverdine is the more potent siderophore.

**Figure 1.**
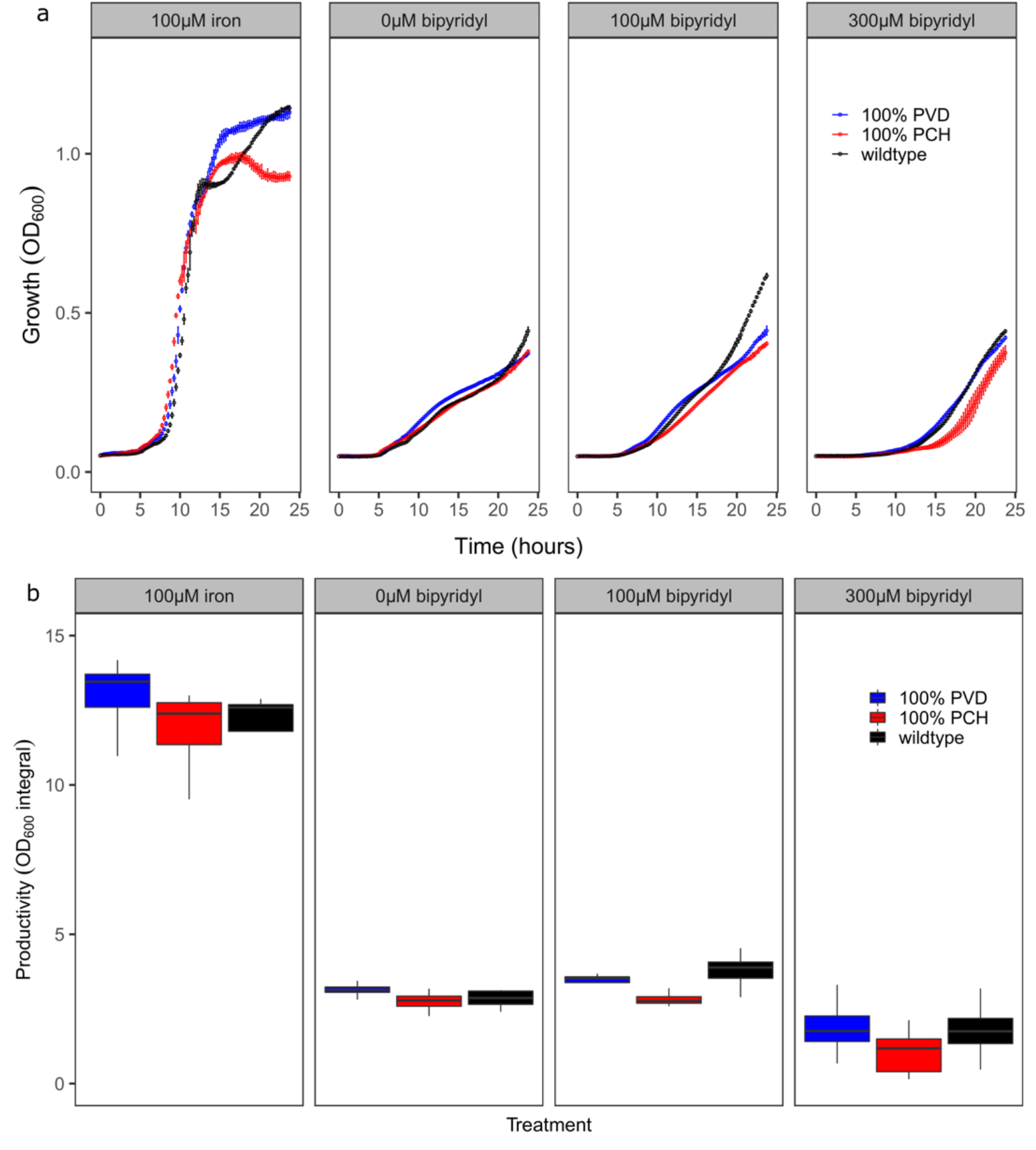
Growth of *P. aeruginosa* strains depend on both iron availability and their siderophore repertoire. Growth and productivity of pyoverdine-producing (blue: *PAO1*Δ*pchEF::mcherry* - 100% PVD) and pyochelin-producing (red: PAO1Δ*pvdD*::egfp - 100% PCH) specialists are compared with the generalist wildtype (black: PAO1), which produces both siderophores in media differing in iron availability. **(a)** Temporal growth dynamics of siderophore specialists and the wildtype. Values and error bars represent mean and standard deviation across 8 replicates, respectively. **(b)** Productivity measured as integral (area under the growth curve, OD_600_) over 24 hours. Boxplots represent the median with 25^th^ and 75^th^ percentiles, and whiskers show the 1.5 interquartile range. Productivity of the pyochelin specialist is significantly lower than the productivity of the pyoverdine specialist (Table S2).

### Specialisation increases productivity in iron rich but not in iron limited media

Next, we investigated how the productivity of mixtures of the two specialist siderophore producers varies across mixing ratios and media conditions, and whether there are conditions under which specialists grow better than the generalist wildtype, which could reflect division of labour. For this purpose, we expressed the productivity of the specialist mixtures relative to the wildtype monoculture. First, we compared productivity between treatments and observed that the productivity significantly declined as the starting frequency of the pyochelin producers in the mixes increased (ANCOVA: F_1,286_= 52.25, p < 0.0001, Table 1) and the iron availability in the medium decreased (F_3,286_= 83.77, p < 0.0001, Table 1). The decline in productivity with more pyochelin specialists in the mix was steeper under more stringent iron limitation (significant interaction between the two variables: F_3,286_= 26.58, p < 0.0001, Table 1). Specifically, mixing ratio did not influence productivity in iron-rich CAA medium (slope: 0.0002, F_1,72_= 0.986, p = 0.3239, Table S3), but did increasingly so when iron became more limiting (plain CAA medium: slope = −0.0008, F_1,72_= 13.90, p = 0.0004; CAA+100μM bipyridyl: slope = −0.0011, F_1,72_= 17.41, p < 0.0001; CAA+300μM bipyridyl: slope = −0.0054, F_1,70_= 36.52, p < 0.0001, Table S3). These results clearly demonstrate that the presence of pyochelin specialists compromises group productivity when iron becomes more limiting.

**Table 1.** The effect of initial pyochelin frequency and iron availability on the relative productivity of specialist mixes and the relative fitness of pyochelin producers.

| Linear model | Relative productivity |  |  | Relative fitness |  |  |
| --- | --- | --- | --- | --- | --- | --- |
| Model statistics | Df | F value | p value | Df | F value | p value |
| Pyochelin frequency (*pch*) | 1,286 | 52.25 | < 0.0001 | 1,224 | 89.73 | < 0.0001 |
| Iron availability (*iron*) | 3,286 | 83.77 | < 0.0001 | 3,224 | 14.03 | < 0.0001 |
| *pch* * *iron* | 3,286 | 26.58 | < 0.0001 | 3,224 | 4.091 | 0.0075 |

In a subsequent step, we compared the relative productivity of the specialist mixes to the generalist wildtype performance and found that mixes had significantly higher productivity than the wildtype in iron-supplemented CAA medium (one-sample t-test: *t*_73_=12.17, p < 0.0001), while there was no significant difference in productivity in plain CAA medium (*t*_73_=1.819, p = 0.0730). In contrast, productivity of mixtures was reduced compared to the wildtype in CAA medium supplemented with bipyridyl (100μM: *t*_73_= −12.89, p < 0.0001; 300μM: *t*_71_= −6.745, p < 0.0001). These findings show that the generalist strategy is costly in iron-rich media where siderophores are less important for iron scavenging, and that specialization saves costs associated with residual siderophore production. However, this cost saving does not result into efficient division of labour under more stringent iron limitation (i.e., in media with bipyridyl).

### Specialists socially interact with each other

We then asked whether the performance of the two siderophore specialists within mixed cultures is simply driven by their intrinsic growth rates shown in the specific medium, or whether there is evidence for social interactions between them. To do so, we calculated the expected frequencies of the pyochelin producer (assuming no social interaction with the pyoverdine producer, see methods), and contrasted them to the observed frequencies in the mixtures, where social interactions are possible. We found that the observed frequencies of the pyochelin producer were significantly different from the expected frequencies (Fig. S2, F_1,454_= 89.12, p < 0.0001). Overall, pyochelin producers performed better than expected with their relative performance being influenced by both their starting frequency (F_1,454_= 40.83, p < 0.0001) and iron availability (F_3,454_= 9.462, p < 0.0001). These results strongly suggest that the two specialists socially interact with one another via siderophore sharing and that this interaction primarily benefits the pyochelin producer.

### Negative-frequency dependent fitness patterns foster specialist co-existence

We then asked whether the two specialists can co-exist or whether one strain would outcompete the other. We thus assessed the relative fitness of the pyochelin producer (relative to the pyoverdine producer) for all mixing ratios. We found that the relative fitness of the pyochelin producer was determined by a significant interaction between mixing ratio and media: F_3,224_= 4.091, p= 0.0075, Fig. 3, Table 1). Specifically, the relative fitness of pyochelin producers significantly declined with higher initial starting frequencies in all media, but the slope of decline differed between media (iron-supplemented CAA: slope = −0.0129, F_1,56_= 73.94, p < 0.0001; plain CAA: slope = −0.0048, F_1,56_= 11.17, p = 0.0015; CAA+100μM bipyridyl: slope = −0.0070, F_1,56_= 21.53, p < 0.0001; CAA+300μM bipyridyl: slope = −0.0075, F_1,56_= 11.27, p = 0.0014, Table S3). Crucially, all regression lines subtend the ln(v) = 0 line, revealing evidence for negative-frequency dependent selection and the possibility of the two specialists to co-exist at an equilibrium. Based on the fitted regression line, we calculated the frequency of pyochelin specialists in the mixes at which a stable equilibrium is reached and found that these values varied across environmental conditions: iron-supplemented CAA: 69.80%; plain CAA: 38.99%; CAA+100μM bipyridyl: 22.29%; CAA+300μM bipyridyl: 30.09%.

## Discussion

In our study we tested whether enforced specialisation in the production of public goods can lead to division of labour. We used the siderophores pyochelin and pyoverdine in the bacterium *P. aeruginosa* as our model system. We mixed specialist strains producing either of the two siderophores at different starting frequencies and subjected them to a gradient of iron limitations. The productivity of the mixes was compared to that of the generalist wildtype and the relative frequency of the specialist were quantified after a 24-hour growth cycle. We found that (i) specialisation increased group productivity under iron-rich conditions where siderophores are not important, (ii) specialist and wildtype cultures performed equally well under moderate iron limitation where both pyochelin and pyoverdine are useful, (iii) specialization reduced group productivity in the environments where iron availability was low and pyoverdine is the more important siderophore, (iv) strains socially interacted with each other, and lastly (v) specialists co-existed in all environments settling on an intermediate ratio. Our observations indicate that enforced specialisation does not lead to an efficient division of labour but fosters coexistence through negative-frequency dependent selection based on mutual cheating under most conditions.

We found that mixtures of the two specialists performed better than the generalist wildtype in iron rich environment (CAA + 100μM iron). However, this finding does not indicate division of labour because siderophores are not important for iron scavenging under such iron-replete conditions (Ochsner *et al*., 1995; Serino *et al*., 1997; Dumas *et al*., 2013). Instead, our results suggest that being a generalist is costly under iron-replete conditions. This interpretation is supported by recent findings at the single cell level showing that the wildtype expresses both pyochelin and pyoverdine synthesis genes at background levels even in iron-rich environments (Mridha & Kümmerli, 2021). Consequently, specialist communities are able to minimize these residual metabolic costs, while the generalist cannot.

We further found that specialist communities were as productive as the generalist wildtype cultures in a medium where the overall iron availability is low, but iron is not bound to a chelator (plain CAA). This observation matches our prediction that division of labour is most likely to arise in environments where both siderophores are beneficial. Pyochelin is beneficial because it is relatively cheap to produce yet is still efficient in iron scavenging despite its relatively low iron affinity (K_f_ = 10^5^ M^-1^) (Youard *et al*., 2011). Pyoverdine is beneficial in any case due to its high iron affinity (K_f_ = 10^24^ M^-1^) (Visca *et al*., 2007), but is produced at moderate amounts presumably because of its relatively high production costs (Dumas *et al*., 2013). Our data therefore suggest that the two iron-scavenging strategies successfully complement each other under moderate iron limitation. However, we might ask why the two specialists only reach wildtype productivity and do not engage in a more efficient division of labour? One explanation is that our strains are imperfect specialist. Pyoverdine and pyochelin synthesis occurs via non-ribosomal peptide synthesis, which involves a series of enzymes (Visca *et al*., 2007; Youard *et al*., 2011). While our mutants lack parts of the siderophore synthesis machinery, which turns them into phenotypic specialists, molecular studies have shown that the remaining part of the synthesis machinery is still partially active resulting into residual metabolic costs (Tiburzi *et al*., 2008). We propose that naturally evolving specialists might alleviate these costs, which could indeed give rise to division of labour under moderate iron limitation.

In contrast, we observed that specialist communities had significantly lower productivity relative to the generalist wildtype cultures in media where iron was bound to a strong iron chelator (CAA + bipyridyl). Under strong iron limitation the importance of pyochelin diminishes, while the one of pyoverdine increases (Dumas *et al*., 2013). This disbalance in their relevance might prevent division of labour, as is demonstrated by our results that community productivity sharply declines with higher initial frequencies of pyochelin specialists in the mixes, showing that they are a burden for the community. Altogether, our productivity analyses show that the conditions under which division of labour, involving siderophore sharing, can evolve is narrow and restricted to environments with moderate iron limitation.

A key prerequisite for division of labour to occur is that specialists interact, and in our case exchange the two siderophores at the community level. Our results support this view as we found that the relative fitness (Fig. 3) and the overall performance (Fig. S2) of each specialist was influenced by the presence and frequency of the other specialist in the community. These findings corroborate previous results that both pyoverdine and pyochelin can be shared between cells and confer fitness benefits to the receivers (Griffin *et al*., 2004; Ross-Gillespie *et al*., 2015; Sathe *et al*., 2019). But albeit the sharing of the two siderophores, enforced specialisation in our system did not result in division of labour but rather stimulated cheating. We reason that this is because the two siderophores have different costs and benefits, especially under strong iron limitation. While pyochelin is relatively cheap to produce, it comes with little benefits for the community when iron is bound to a strong chelator (see also Fig. 1). Conversely, pyoverdine is relatively costly for the producers yet generates high benefits for the community as it manages to scavenge iron even when it is bound to a strong chelator. This imbalance in costs and benefits puts pyochelin producers at an advantage as they can exploit the beneficial pyoverdine while saving costs by making the cheaper (but less useful) pyochelin. Consequently, pyochelin producers act predominantly as cheater under severe iron limitation, reaping relative fitness benefits (Fig. 3) while dragging community productivity down (Fig. 2). But our data suggests that pyoverdine producers can also cheat on pyochelin producers, especially when rare (Fig. S3). At low pyoverdine producer frequencies, the balance tips and our analysis shows that pyochelin producers reach lower frequencies than expected based on their monoculture growth, both under moderate (plain CAA) and strong (CAA + bipyridyl) iron limitation (Fig. S3). Taken together, these results suggest that interactions between specialists are characterized by mutual cheating and not division of labour under iron limitation.

**Figure 2.**
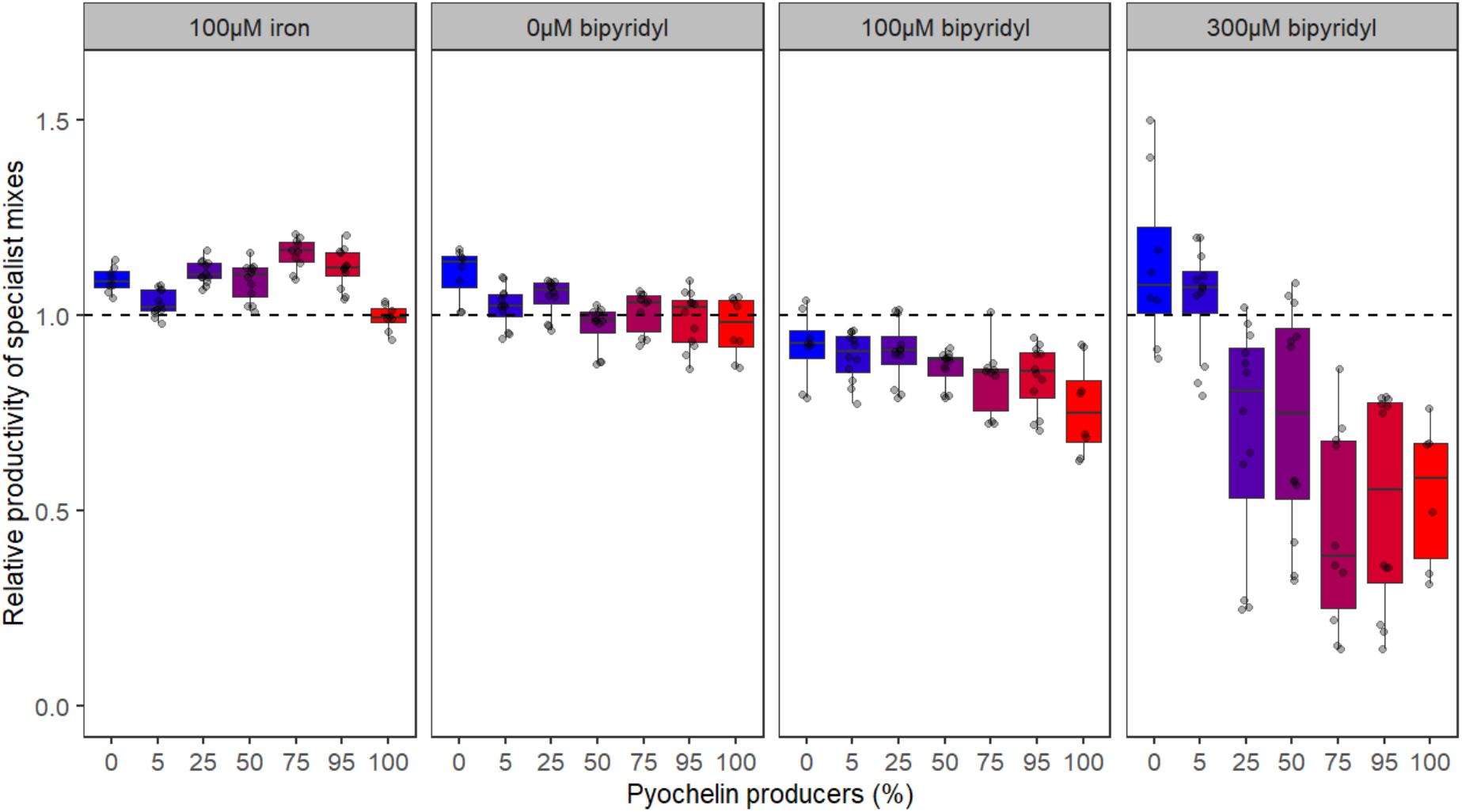
Siderophore specialization increases productivity in iron-rich medium, while it declines with higher starting frequencies of pyochelin producers in iron-limited media. The colour code depicts mixing ratios from 0% pyochelin producers (pyoverdine monoculture) in blue to 100% pyochelin producers (pyochelin monoculture) in red, with the increase in frequency of pyochelin producers in the mixes being denoted by a gradual shift from blue to red. Relative productivity is measured as the growth integral (area under the OD_600_ curve) of the specialist monocultures and mixes (*PAO1*Δ*pvdD::egfp* + PAO1Δ*pchEF::mcherry*) divided by the growth integral of the wildtype (PAO1). The dashed line indicates the relative productivity of the wildtype. Boxplots represent the median with 25^th^ and 75^th^ percentiles, and whiskers show the 1.5 interquartile range across 12 replicates.

**Figure 3.**
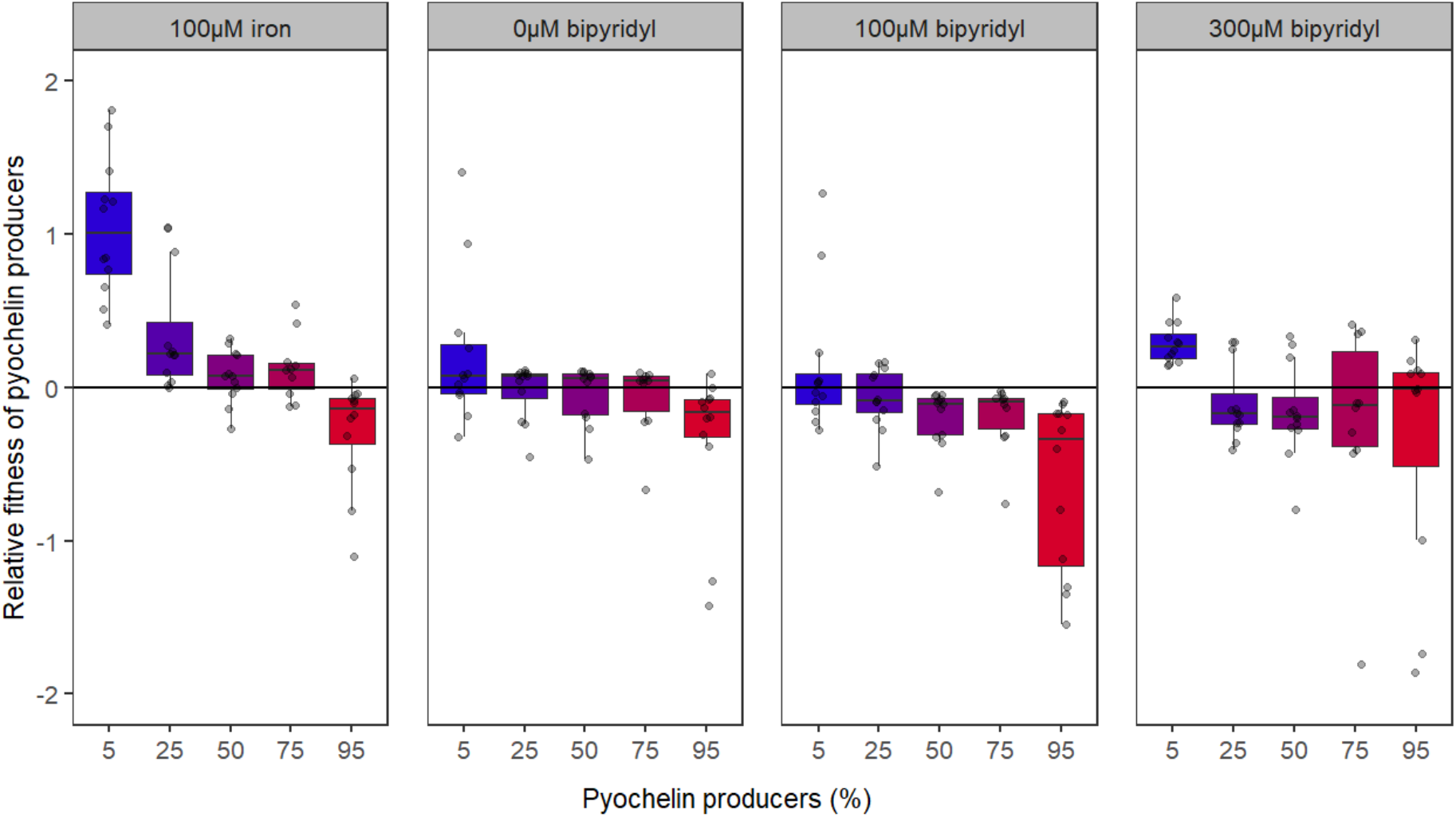
Relative fitness ln(*v*) of pyochelin producers in competition with pyoverdine producers declines with higher starting frequencies of pyochelin producers in the population. The colour code depicts mixing ratios from 0% pyochelin producers (pyoverdine monoculture) in blue to 100% pyochelin producers (pyochelin monoculture) in red, with the increase in frequency of pyochelin producers in the mixes being denoted by a gradual shift from blue to red. These findings suggest that there is negative-frequency dependent selection in all media conditions. Competitions occurred between pyochelin (*PAO1*Δ*pvdD::egfp*) and pyoverdine (PAO1Δ*pchEF::mcherry*) producers over 24 hours in media differing in iron availability. Values of ln(*v*) > 0, ln(v) = 0, or ln (*v*) <0 indicate that pyochelin producers won, did equally well or lost in competition against pyoverdine producers, respectively. Boxplots represent the median with 25^th^ and 75^th^ percentiles, and whiskers show the 1.5 interquartile range across 12 replicates.

Yet despite cheating, we observed co-existence between the two specialists through negative frequency dependent selection under all conditions. The question is whether co-existence would be stable in the long run? One conceivable option is that a double mutant would arise that cheats on both siderophores, potentially bringing the communities to collapse under strong iron limitation. However, a previous experimental evolution study yielded no evidence for such a scenario (Ross-Gillespie *et al*., 2015), but showed stable co-existence of the wildtype generalist with a pyoverdine mutant that overproduced pyochelin and a second mutant that produced both siderophores at intermediate levels. An alternative option is that the interactions between specialists is lost through the evolution of an independent lifestyle, as shown to occur in the enforced biofilm specialization system in *Bacillus subtilis* (Dragoš *et al*., 2018b). In our case, this could happen through the evolution of a superior siderophore system that is neither shareable nor exploitable by other community members (Lee *et al*., 2012).

We showed that enforced specialization saves metabolic costs associated with the production of the two siderophores (Fig. 2, iron-rich conditions). However, this cost saving did not result in an efficient division of labour under iron-limited conditions. We propose that this is because *P. aeruginosa* has evolved sophisticated regulatory mechanisms to fine-tune siderophore investment in response to environmental conditions. Such regulatory mechanisms also save costs (Kümmerli and Brown, 2010; Xavier *et al*., 2011; Darch *et al*., 2012) and could represent an alternative adaptive strategy to division of labour. Siderophore regulation in *P. aeruginosa* involves three levels. At the top level, Fur (ferric uptake regulator) blocks siderophore synthesis when iron accumulates within cells but loses its inhibitory effect when intra-cellular iron reserves become depleted (Leoni *et al*., 1996; Ochsner, 1996; Escolar *et al*., 1999; Youard *et al*., 2011). Fur repression seems to act more strongly on pyoverdine than pyochelin, allowing for preferential pyochelin production under moderate iron limitation (Dumas *et al*., 2013). The second level of regulation involves membrane-embedded signalling cascades, where incoming siderophore-iron complexes trigger a positive feedback loop that increases siderophore production in response to successful scavenging events (Lamont *et al*., 2002; Michel *et al*., 2007). The third level is poorly explored at the molecular level, yet is hierarchical in nature, whereby pyoverdine production suppresses pyochelin synthesis intracellularly (Dumas *et al*., 2013), allowing for preferential pyoverdine production under strong iron limitation. Such regulatory circuits could outperform genetic division of labour because individuals remain generalists and thus flexible in responding to environmental fluctuations (Butaitė *et al*., 2018), while genetic division of labour requires specialist partners to be present in the environment in adequate ratios and being able to exchange their goods, conditions that might often not be met.

In conclusion, our study indicates that enforcing specialisation with regard to siderophore production does not lead to beneficial division of labour in *P. aeruginosa* but leads to the stable co-existence of the two specialists through mutual cheating. Our results suggest that there are at least three factors that could constrain the evolution of division of labour with regard to public good exchange in bacterial population. First, an imbalance in the costs and benefits of two public goods can promote cheating rather than an efficient division of labour. Second, division of labour might only be beneficial under specific environmental conditions and because environmental fluctuations are common in natural habitats generalist strategies might be favoured. Third, the evolution of fine-tuned regulatory circuits can compensate for the costs associated with pursuing a generalist strategy, and at the same time allows generalist to flexibly respond to environmental fluctuations. Given these constraints, genetic division of labour might be harder to evolve in bacteria than previously thought, especially when environmental conditions fluctuate in unpredictable ways, making the evolution of fine-tuned regulatory circuits the better option.

## Acknowledgments

We thank Priyanikha Jayakumar, Selina Niggli, Santosh Sathe and Martina Archetti for help in the lab and with the data analysis, and the Cytometry Facility at University of Zurich. This project has received funding from the European Research Council (ERC) under the European Union’s Horizon 2020 research and innovation programme (grant agreement no. 681295).

## Author contributions

SM and RK designed the study. SM carried out all experiments. SM and RK analysed the data and wrote the paper.

