## Supplementary Material for "Enforced specialization fosters mutual cheating and not division of labour in the bacterium *Pseudomonas aeruginosa*"

This file contains the following supplementary materials:

- 3 supplementary figures
- 3 supplementary tables

### Supplementary figures

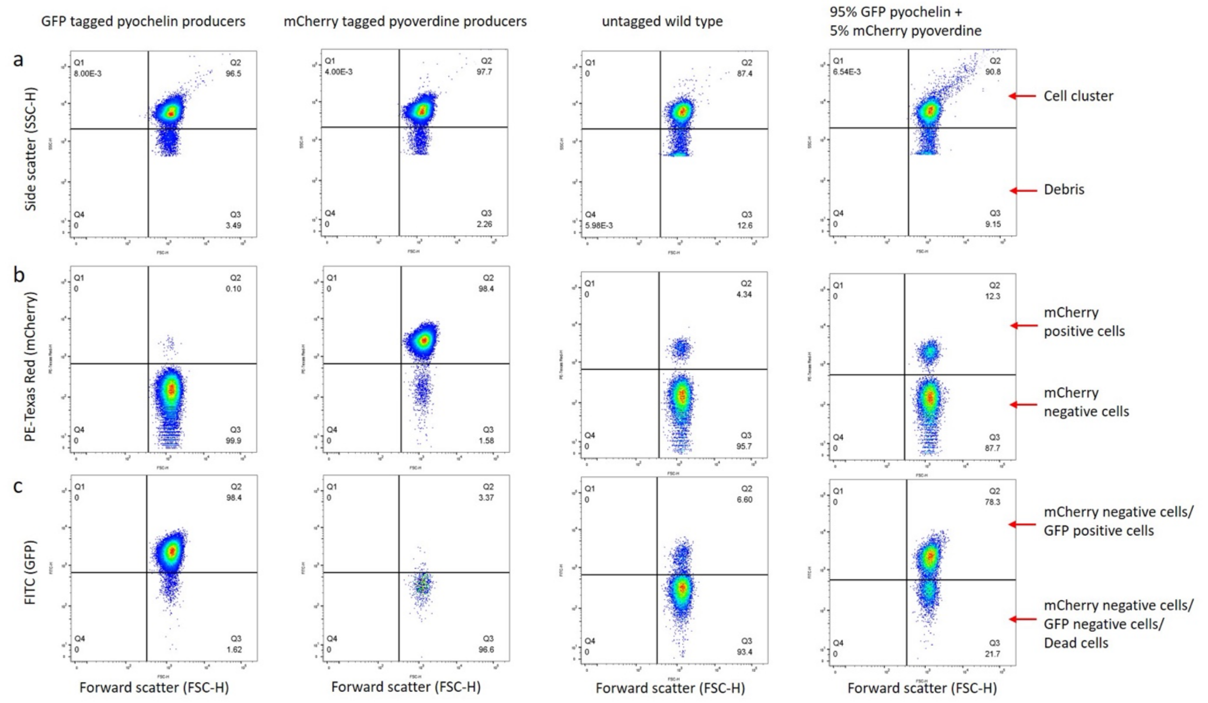


**Figure S1.** Gating strategy for flow cytometry data, explaining how we distinguished and quantified the final frequency of mCherry tagged (PAO1Δ*pchEF::mcherry*) pyoverdine producers and GFP tagged (PAO1Δ*pvdD::gfp*) pyochelin producers after 24 hours of growth across media differing in iron availability. GFP tagged pyochelin and mCherry tagged pyoverdine producer monocultures were used as positive control. Untagged wild type monoculture was used as negative control and a mix containing 95% pyochelin and 5% pyoverdine producers is shown as a test case. **(a)** Four panels in top row; we separated the cells from debris, by using the forward (FSC-H) and side scatter (SSC-H) as a proxy for particle size. **(b)** Four panels in middle row; within that cell cluster we imposed a PE-Texas Red-H (mCherry) and forward scatter (FSC-H) gate to distinguish between mCherry positive (pyoverdine producers) and negative cells**.** **(c)** Four panels in bottom row; within the mCherry negative cell cluster, we then imposed the FITC-H (GFP) and forward scatter (FSC-H) gates to distinguish between GFP positive (pyochelin producers) and negative cells (dead cells). The mCherry positive and GFP positive cells were then recorded and their final frequencies were calculated after accounting for the debris and dead cells.

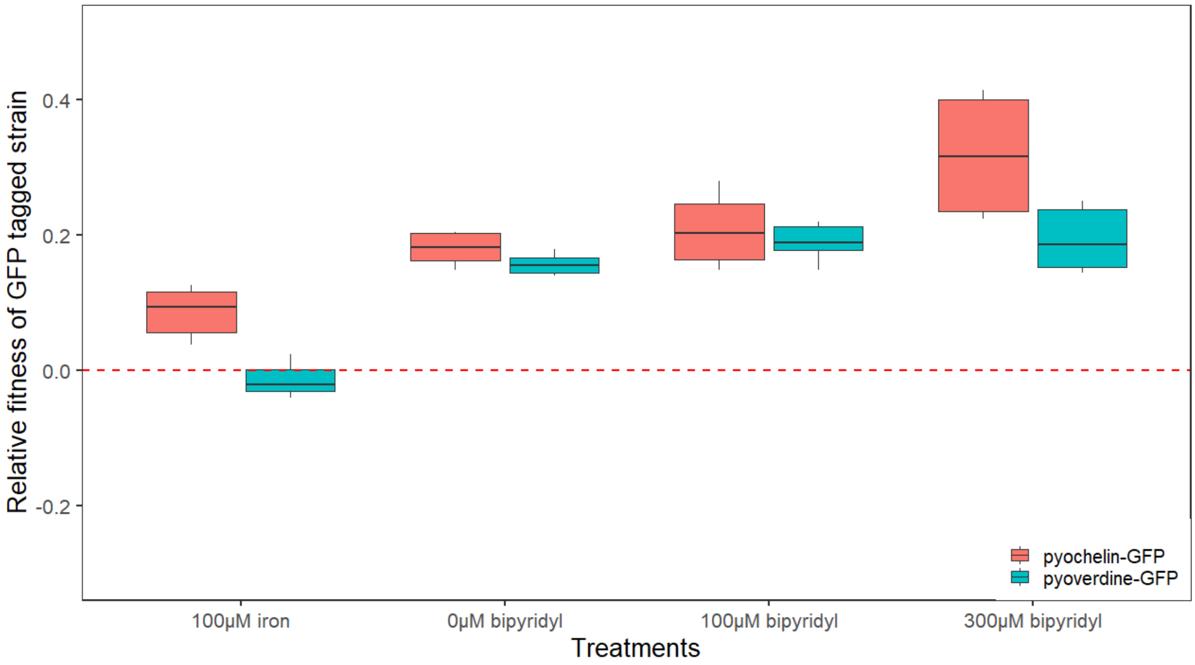


**Figure S2.** Relative fitness ln(*v)* of GFP expressing siderophore producers in competition with its identical mCherry expressing variant in 1:1 ratio across media differing in iron availability. GFP expressing siderophore producers outcompetes mCherry expressing variants. Boxplots represent the median with $25^{th}$ and $75^{th}$ percentiles, and whiskers show the 1.5 interquartile range across 6 replicates.

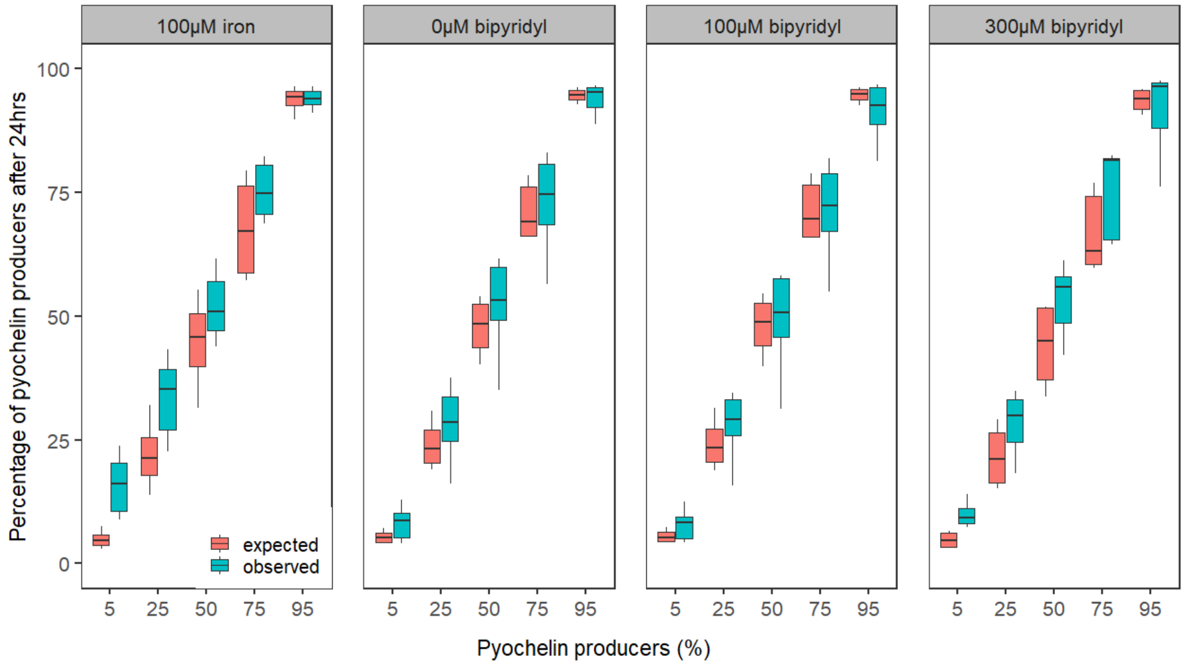


**Figure S3.** Pyochelin producers grow better than expected in mixed cultures. The observed final frequency was obtained (with flow cytometry) as a percentage of pyochelin producers after 24-hour growth in media differing in iron availability. The expected frequency is estimated based on the intrinsic growth rate and time spent in the exponential phase by pyochelin producers (PAO1Δ*pvdD::gfp*) and pyoverdine producers (PAO1Δ*pchEF::mcherry*). Boxplots represent the median with $25^{th}$ and $75^{th}$ percentiles, and whiskers show the 1.5 interquartile range.

**Table S1.** Average relative fitness of specialist GFP tagged siderophore producers in competition with specialist mCherry tagged siderophore producers.

| Medium | Average relative fitness of GFP tagged strain |
|---|---|
| 100 μM iron | 0.0359 |
| 0 μM bipyridyl | 0.1641 |
| 100 μM bipyridyl | 0.1981 |
| 300 μM bipyridyl | 0.2654 |

**Table S2.** Monocultures growth over 24 hours across a range of CAA medium differing in their iron availability.

| **Strain** | **Medium** | **Integral_OD600** | **MaxSlope_Growthrate** |
|---|---|---|---|
| 100% PVD | 100µM iron | 12.95328923 | 0.220310075 |
| 100% PVD | 0µM bipyridyl | 3.123454675 | 0.022895508 |
| 100% PVD | 100µM bipyridyl | 3.437661763 | 0.032142723 |
| 100% PVD | 300µM bipyridyl | 1.883748575 | 0.045415435 |
| 100% PCH | 100µM iron | 11.82288555 | 0.227744575 |
| 100% PCH | 0µM bipyridyl | 2.745419363 | 0.01911874 |
| 100% PCH | 100µM bipyridyl | 2.8306653 | 0.031804473 |
| 100% PCH | 300µM bipyridyl | 1.062779567 | 0.033693926 |
| wildtype | 100µM iron | 11.8952764 | 0.229599539 |
| wildtype | 0µM bipyridyl | 2.82954955 | 0.040708183 |
| wildtype | 100µM bipyridyl | 3.780997488 | 0.275169018 |
| wildtype | 300µM bipyridyl | 1.783981513 | 0.044682769 |

Table S3. The effect of initial pyochelin frequency on the relative productivity of the specialist mixes and relative fitness of pyochelin producers in the four environments differing in iron availability.

| Linear model | Relative productivity | | | | Relative fitness | | | |
|---|---|---|---|---|---|---|---|---|
| Model statistics | Df | F value | P value | Slope | Df | F value | P value | Slope |
| CAA + 100µM iron | 1,72 | 0.986 | 0.3239 | 0.0002 | 1,56 | 73.94 | < 0.0001 | -0.0129 |
| plain CAA medium | 1,72 | 13.90 | 0.0004 | -0.0008 | 1,56 | 11.17 | 0.0015 | -0.0048 |
| CAA + 100µM bipyridyl | 1,72 | 17.41 | 0.0001 | -0.0011 | 1,56 | 21.53 | < 0.0001 | -0.0070 |
| CAA + 300µM bipyridyl | 1,70 | 36.52 | 0.0001 | -0.0054 | 1,56 | 11.27 | 0.0014 | -0.0075 |
